# Evolutionary genomics identifies host-directed therapeutics to treat intracellular bacterial infections

**DOI:** 10.1101/2023.08.01.551011

**Authors:** Josette Medicielo, Eric Waltari, Abigail Leigh Glascock, Gytis Dudas, Brian DeFelice, Ira Gray, Cristina M. Tato, Joan Wong, Vida Ahyong

## Abstract

Obligate intracellular bacteria shed essential biosynthetic pathways during their evolution towards host dependency, providing an opportunity for host-directed therapeutics. Using *Rickettsiaceae* as a model, we employed a novel computational pipeline called PoMeLo to systematically compare this cytosolic family of bacteria to the related *Anaplasmataceae*, which reside in a membrane-bound vacuole in the host cell. We identified 20 metabolic pathways that have been lost since the divergence of *Anaplasmataceae* and *Rickettsiaceae*, corresponding to the latter’s change to a cytosolic niche. We hypothesized that drug inhibition of these host metabolic pathways would reduce the levels of metabolites available to the bacteria, thereby inhibiting bacterial growth. We tested 22 commercially available inhibitors for 14 of the identified pathways and found that the majority (59%) reduced bacterial growth at concentrations that did not induce host cell cytotoxicity. Of these, 5 inhibitors with an IC_50_ under 5 μM were tested to determine whether their mode of inhibition was bactericidal or bacteriostatic. Both mycophenolate mofetil, an inhibitor of inosine-5’-monophosphate dehydrogenase in the purine biosynthesis pathway, and roseoflavin, an analog of riboflavin, displayed bactericidal activity. A complementary unbiased mass spectrometry-based metabolomics approach identified 14 pathways impacted by *Rickettsia* infection based on alterations in metabolite levels. Strikingly, 11 of these (79%) overlapped with those identified by our computational predictions. These *in vitro* validation studies support the feasibility of a novel evolutionary genomics-guided approach for host-directed antibiotic drug development against obligate pathogens.

**Importance:** Many pathogens have evolved to acquire essential metabolites from their host cell, while in turn shedding their own biosynthetic capacities. This leads to an interesting dilemma: on one hand, reduced genomes allow pathogens to save energy and replicate more quickly, while on the other hand, they become more dependent on the host cell for survival. This vulnerability can be exploited by identifying and therapeutically inhibiting the host pathways that are essential for pathogen survival. The significance of our research is in predicting the precise pathways lost during a pathogen’s evolutionary adaptation to parasitism and validating these predictions through targeted *in vitro* growth assays and an unbiased metabolomic survey of the host-pathogen interface.

## Introduction

As microbes evolve to become more associated with their hosts, some make the switch from free-living organisms to obligate pathogens, becoming fully dependent on the host cell for growth. Previous studies have described the phenomenon of genomic streamlining in pathogen and symbiont genomes over long evolutionary timescales, during which genes and biosynthetic pathways that typically allow for a free-living lifestyle are shed (Andersson and Andersson, 1999; Merhej et al., 2013; Moran, 2002; Wolf and Koonin, 2013). Understanding how these lost pathways in pathogen genomes underlie host dependency could lead to novel avenues for antimicrobial interventions.

Gram-negative, obligate, intracellular bacteria from the Order Rickettsiales have undergone massive genome streamlining, with genome sizes generally smaller than 2 Mb (Andersson and Andersson, 1999; Andersson et al., 1998; Blanc et al., 2007; Darby et al., 2007; Merhej et al., 2013; Merhej and Raoult, 2011). The Order Rickettsiales is further subdivided into two major families: the *Anaplasmataceae* and the *Rickettsiaceae* (Darby et al., 2007; Salje, 2021; Yu and Walker, 2006). The *Anaplasmataceae* family includes bacteria from the genera *Wolbachia, Anaplasma, Neorickettsia*, and *Ehrlichia* that have been classified as either parasitic or symbiotic with their host. The *Rickettsiaceae* family include the genera *Orientia* and *Rickettsia* and are largely considered to be pathogens of both humans and animals. One fundamental difference between the two families is their intracellular growth niche, with the *Anaplasmataceae* residing within a membrane-bound vacuole, and the *Rickettsiaceae* escaping from the vacuole and residing and replicating in the host cytoplasm (Hackstadt, 1996). This key distinction could further influence the evolutionary and genomic relationship between the intracellular bacteria and their hosts.

Previous studies have explored host metabolic contribution to rickettsial metabolism, using phylogenomic and computational approaches to identify missing metabolic capabilities within the rickettsial genome (Driscoll et al., 2017; Fuxelius et al., 2007). Both studies concluded that metabolic gaps are functionally fulfilled by parasitizing host metabolites. The host terpenoid pathway was identified in the Driscoll *et al*. study as essential for the synthesis of bacterial ubiquinone and lipid carriers for peptidoglycan and lipopolysaccharides. Our previous work posited that in the *Rickettsiaceae*, the decay of the terpenoid pathway was likely enabled by its cytoplasmic access to host terpenoid precursors, unlike the vacuole-bound *Anaplasmataceae* that still encode the bacterial terpenoid pathway (Ahyong et al., 2019). We tested the hypothesis that *Rickettsia parkeri* acquires host terpenoid precursors and found that *in vitro* inhibition of the host terpenoid pathway with statins inhibits bacterial growth. This finding leads us to ask whether a viable option for antimicrobial therapy would be to suppress production of additional pathogen-enabling metabolites in the host.

Traditionally, antimicrobial agents target pathogen-specific proteins or processes. However, the paucity of new antibiotics being developed suggests that this approach is no longer as fruitful (Kaufmann et al., 2018). Furthermore, the rapid generation of drug resistance due to a pathogen’s ability to degrade, efflux, or mutate the target protein of an antibiotic drug is problematic for future drug discovery efforts. One promising approach that avoids the complications of pathogen-targeted therapeutics while expanding the number of potential target proteins and processes is host-directed therapeutics, which has been proposed for viral and *Mycobacterium tuberculosis* infections (Hawn et al., 2013; Kaufmann et al., 2018; Zumla et al., 2016). Host-directed therapeutic approaches differ in their mechanisms of pathogen growth inhibition. Some examples include modulating the host immune response to pathogen detection or clearance, blocking access of essential host proteins that initiate pathogen attachment or entry, and reducing the pathophysiological damage to the host. Furthermore, drug resistance to a host-directed therapeutic is unlikely to generate resistant microbes because the therapeutic target is not encoded within the pathogen genome. Therefore, we explored whether host metabolic pathways could serve as promising targets for antimicrobial intervention, focusing specifically on the evolutionary changes that occurred when the ancestor of *Rickettsiaceae* escaped the vacuole and entered the nutrient-rich host cytoplasm.

## Results

### Genome streamlining in the Rickettsiales and escape from the vacuole

To systematically identify phylogenetic lineages that have undergone genome reduction, we compared two closely related families (the *Rickettsiaceae* and *Anaplasmataceae*) that include obligate intracellular pathogens and symbionts. To contextualize the extent of genome size reduction in the Rickettsiales in terms of their phylogenetic relationship to other Alphaproteobacteria, we selected a representative set of Alphaproteobacteria genomes (one per genus) and all Rickettsiales genomes that are marked as complete in NCBI’s Bacterial and Viral Bioinformatics Resource Center (BV-BRC) (n=122 for the *Rickettsiaceae* and n=150 for the *Anaplasmataceae*; Supplemental File 1, 2). Using BV-BRC, we extracted genome size information and generated a phylogenetic tree using the “Bacterial phylogenetic tree” tool (Olson et al., 2023). To quantify genome streamlining, we used phylogenetically weighted mean genome sizes to estimate the changes in genome size across the Alphaproteobacteria phylogeny (Figure 1, see Methods). We found that all Rickettsiales have genomes sizes well below the 95th percentile range of their closest Alphaproteobacteria relatives (Figure 1), with further streamlining events occurring within *Anaplasmataceae*, particularly in the genera *Anaplasma, Wolbachia*, and *Ehrlichia* (<400 kb lost) and the genus *Neorickettsia* (>800 kb lost). Unlike the *Anaplasmataceae*, however, members of the *Rickettsiaceae* (genera *Orientia* and *Rickettsia* in Figure 1) have developed the capability to inhabit the nutrient-rich cytoplasm of the host cell. Therefore, we hypothesized that the *Rickettsiaceae* underwent more severe metabolic pathway reduction, in concordance with their reliance on direct siphoning of metabolites from the host cytoplasm.

**Figure 1:**
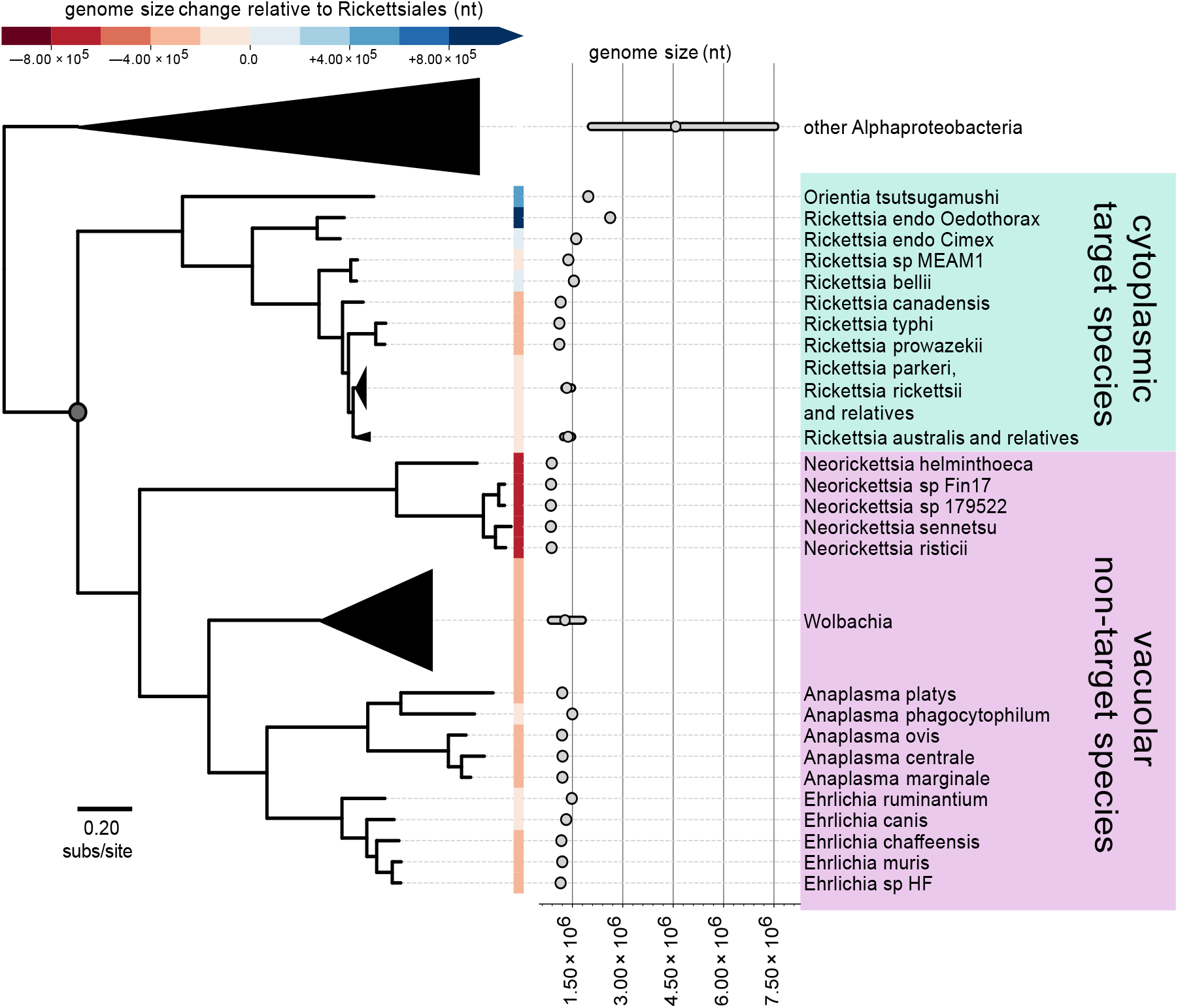
Genomic reduction in the Rickettsiales. A phylogenetic reconstruction of two major families in the Rickettsiales, the *Rickettsiaceae* and the *Anaplasmataceae*, and representative Alphaproteobacteria. Phylogenetically weighted means were computed to quantify genome size differences (in nucleotides) along a phylogenetic tree relative to the Rickettsiales.

### A computational pipeline for identifying metabolic pathway gaps & *predicting host reliance*

We developed a computational pipeline called PoMeLo (Predictor of Metabolic Loss) that can predict the loss of metabolic capacity in a **target** pathogen group, the *Rickettsiaceae*, as compared to a **non-target** group, the *Anaplasmataceae*. The details of the pipeline are described elsewhere (Glascock, Waltari *et al*, bioRxiv 2023). Briefly, protein annotation files for each species are sourced from the BV-BRC and are used to record the presence of genes within metabolic pathways across all species. We first determine the percentage of species that encode each enzyme within the target and non-target group. The percent differential between the groups is then calculated and summed across all enzymes in a pathway to obtain a final Predicted Metabolic Loss (PML) score, where a higher PML score implies a greater loss in metabolic capacity.

Finally, a ranked list of PML scores is reported across 165 metabolic pathways, and heatmap visualizations are provided to further interrogate gene and pathway loss across the target and non-target comparison groups (Figure 2a; Supplemental File 3).

**Figure 2:**
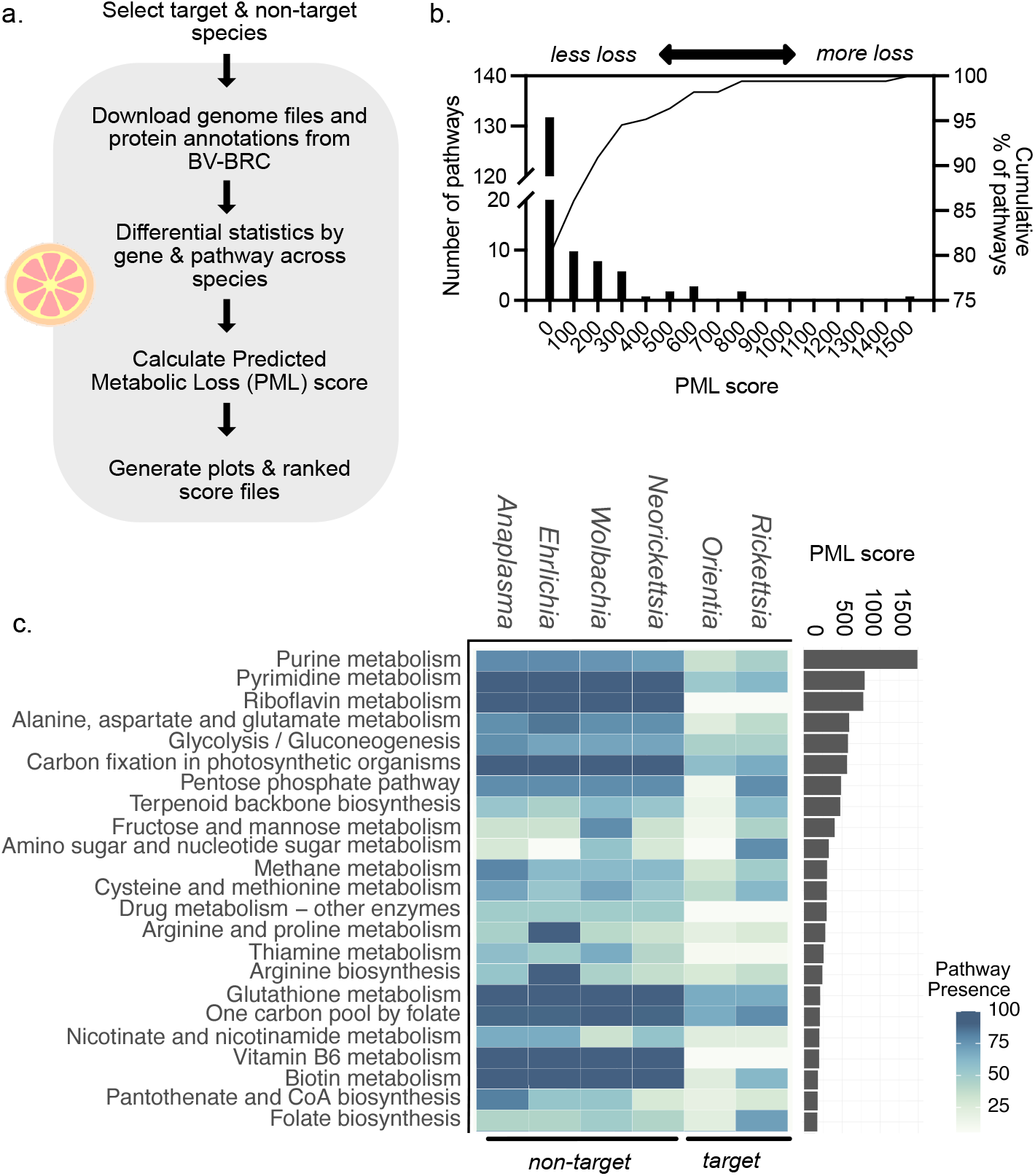
PoMeLo pipeline predicts missing metabolic pathways in the target *Rickettsiaceae* group compared to the non-target *Anaplasmataceae* group. a) Schematic of the PoMeLo workflow. b) Distribution of PML scores for all metabolic pathways analyzed in the target versus non-target comparison. c) Heatmap showing the overall pathway presence percentage and PML scores for the top 23 pathways ranked by PoMeLo across the 6 main genera of the Rickettsiales.

### Metabolic pathway analysis

We predict that the pathways with the highest PML scores (i.e., those coinciding with the gain in ability of the *Rickettsiaceae* to live within the cytoplasm) would have the greatest loss in metabolic capacity and therefore the highest dependence on host metabolites. A histogram of the PML scores for all pathways in the *Rickettsiaceae* versus *Anaplasmataceae* comparison is displayed in Figure 2b. Most pathways show little difference (139 pathways with a PML score ≤ 150). However, 23 pathways (16% of total) displayed gene loss specifically within the target group (Table 1). A heatmap that plots the genus-level pathway presence versus the PML score illustrates the difference in metabolic capacity for the top-ranking pathways selectively lost in the *Rickettsiaceae* (Figure 2c).

**Table 1:** Top 23 pathways scored by the PoMeLo pipeline. The percent of enzymes missing in the target and non-target groups are an average across all species in the respective group.

| Pathway name | PoMeLo Rank | PML score | Percent missing in target group | Percent missing in non-target group |
| --- | --- | --- | --- | --- |
| Purine metabolism | 1 | 1493 | 53.1 | 18.8 |
| Pyrimidine metabolism | 2 | 800 | 38.1 | 0 |
| Riboflavin metabolism | 3 | 782 | 100 | 0 |
| Alanine, aspartate and glutamate metabolism | 4 | 596 | 61.5 | 7.7 |
| Glycolysis / Gluconeogenesis | 5 | 580 | 30.8 | 15.4 |
| Carbon fixation in photosynthetic organisms | 6 | 568 | 16.7 | 0 |
| Pentose phosphate pathway | 7 | 488 | 22.2 | 22.2 |
| Terpenoid backbone biosynthesis | 8 | 478 | 38.5 | 38.5 |
| Fructose and mannose metabolism | 9 | 404 | 44.4 | 11.1 |
| Amino sugar and nucleotide sugar metabolism | 10 | 327 | 22.2 | 38.9 |
| Methane metabolism | 11 | 304 | 50 | 20 |
| Cysteine and methionine metabolism | 12 | 301 | 38.5 | 23.1 |
| Drug metabolism - other enzymes | 13 | 298 | 50 | 0 |
| Arginine and proline metabolism | 14 | 280 | 73.3 | 0 |
| Thiamine metabolism | 15 | 257 | 83.3 | 0 |
| Arginine biosynthesis | 16 | 241 | 63.6 | 0 |
| Glutathione metabolism | 17 | 212 | 33.3 | 0 |
| One carbon pool by folate | 18 | 208 | 22.2 | 0 |
| Nicotinate and nicotinamide metabolism | 19 | 201 | 55.6 | 22.2 |
| Vitamin B6 metabolism | 20 | 198 | 100 | 0 |
| Biotin metabolism | 21 | 181 | 38.5 | 0 |
| Pantothenate and CoA biosynthesis | 22 | 177 | 72.7 | 9.1 |
| Folate biosynthesis | 23 | 173 | 21.4 | 35.7 |

The highest-ranked PML score was for the purine metabolism pathway (PML=1493). Fourteen enzymes for purine metabolism were detected as being completely absent in the *Rickettsiaceae* but >90% present in the *Anaplasmataceae*, as shown in the species-level heatmap (Figure 3a; heatmaps of all species-level and target vs non target groups are shown in Supplemental Figures 1 and 2, respectively).

**Figure 3:**
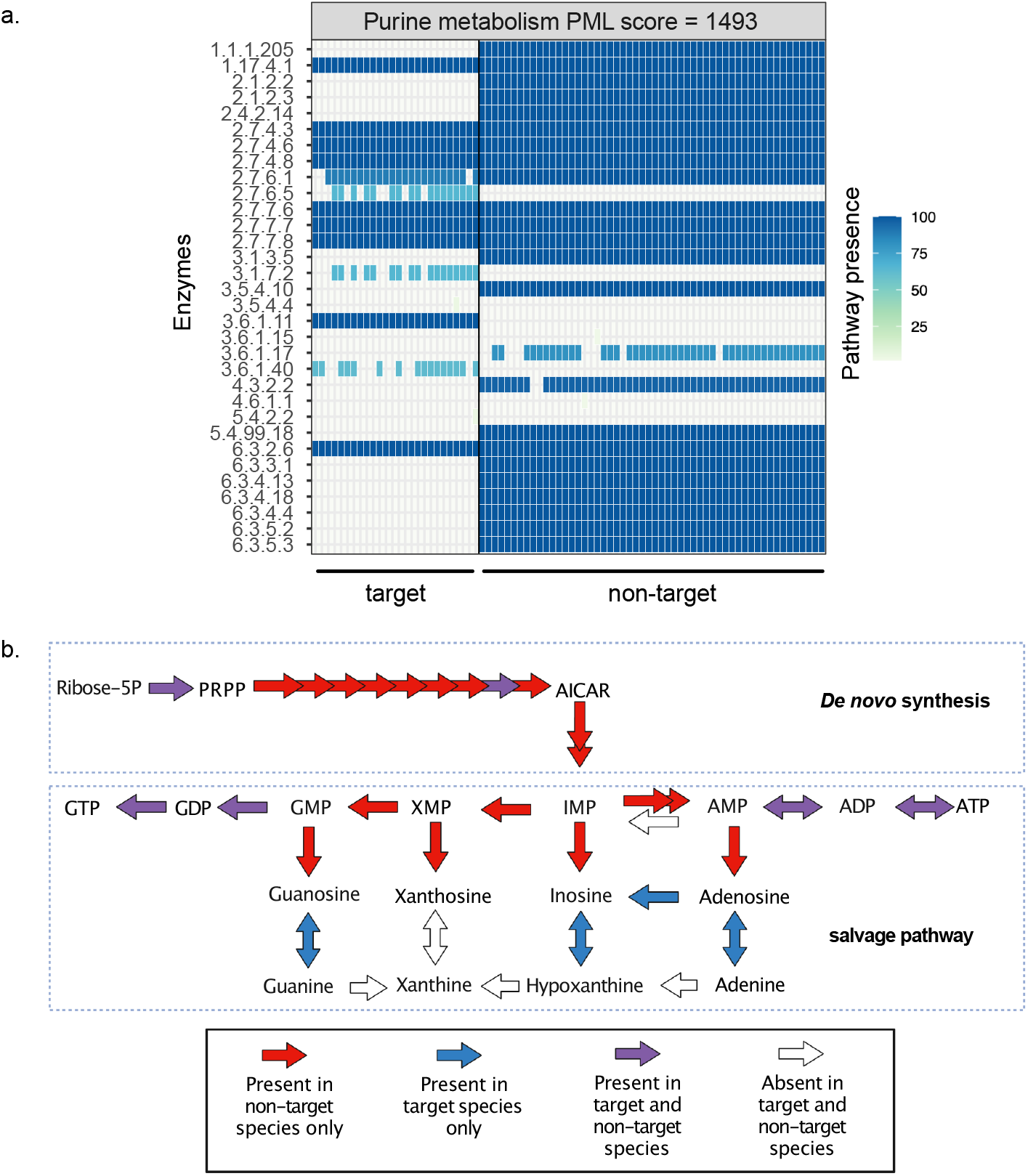
The purine metabolism pathway comparison between the target and non-target groups. a) Heatmap of enzyme presence for target vs non-target species for the purine metabolism pathway. b) Schematic pathway map of the enzyme presence or absence for the purine metabolism pathway, color-coded according to panel A (created using BioRender).

To better visualize where these enzymes occur in the pathway, we mapped each enzyme shown in Figure 3a on a schematic pathway map according to its presence in the target or non-target groups (Figure 3b). While the majority of the enzymes are encoded in the non-target *Anaplasmataceae* group, many are absent in the target *Rickettsiaceae* group. Of the 23 pathways displaying the greatest gene loss, 3 pathways (carbon fixation in photosynthetic organisms, methane metabolism, and drug-metabolism) were excluded from further analysis because they are not likely to play an important role in these bacteria, and many of their missing enzymes were already identified as gaps in other essential pathways (Supplemental Table 1). The final 20 pathways exemplify recent evolutionary reduction or elimination of pathogen-encoded metabolic capacity and a switch to parasitism of host metabolism.

### Validation of host-directed therapeutics against R. parkeri

To test our hypothesis that the pathogens are co-opting host metabolism, we identified inhibitors of host metabolic pathways corresponding to those predicted to be lost in the *Rickettsiaceae*. We obtained 22 commercially available inhibitors for 14 of the 20 pathways selected for *in vitro* susceptibility testing (Table 2). No suitable commercially available inhibitors were identified for the remaining 6 pathways. To assess the efficacy and toxicity of these drugs, we measured the effect of drug concentration on the growth of *R. parkeri-*infected A549 cells, measuring bacterial growth by qPCR of infected cells and host cytotoxicity by lactose dehydrogenase release.

**Table 2:** Drugs tested for each pathway, known drug target, and IC_50_ in μM from the dose response curves.

| Pathway name | Drugs tested | Drug target | IC<sub>50</sub> μM | F= full inhibition, no cytotoxicity<br>C = inhibition + cytotoxicity<br>N = no inhibition<br>P = partial inhibition |
| --- | --- | --- | --- | --- |
| Purine metabolism | Mycophenolate mofetil | IMPDH | 1.13 | F |
|  | Ribavirin | IMPDH | 97.37 | F |
|  | Lometrexol | Glycinamide ribonucleotide formyltransferase | 6.7 | F |
| Pyrimidine metabolism | Brequinar | DHODH | 0.86 | F |
|  | Sparfosic acid | aspartate transcarbamoyl transferase | 133.8 | C |
| Riboflavin metabolism | Lumiflavine | Riboflavin uptake inhibitor | 457.4 | F |
|  | Roseoflavin | Riboflavin uptake inhibitor | 0.6 | F |
| Alanine, aspartate and glutamate metabolism | Telaglenastat | Glutaminase | 5.43 | P |
| Glycolysis / Gluconeogenesis | AP-III-a4 hydrochloride | Enolase | 26.33 | C |
|  | 3PO | 6-phosphofructo-2-kinase/fructose-2,6-biphosphatase 3 | 142.4 | C |
|  | Sodium dichloracetate | Pyruvate dehydrogenase kinase | 2758 | C |
|  | CBR-470-1 | phosphoglycerate kinase 1 | 38.5 | C |
| Carbon fixation in photosynthetic organisms | excluded | NA |  |  |
| Pentose phosphate pathway | 6-amininicotinamide | 6-phosphogluconate dehydrogenase | 1.16 | F |
| Terpenoid backbone biosynthesis | Pitavastatin | HMG-CoA reductase | 4.45 | F |
| Fructose and mannose metabolism | FBPase inhibitor | Fructose-1,6-bisphosphatase-1 | 42.25 | F |
| Amino sugar and nucleotide sugar metabolism | not tested | NA |  |  |
| Methane metabolism | excluded | NA |  |  |
| Cysteine and methionine metabolism | DL-propargylglycine | cystathionine γ-lyase | | F |
| Drug metabolism - other enzymes | excluded | NA |  |  |
| Arginine and proline metabolism | Tetrahydro-2-furoic acid | proline dehydrogenase | NA | N |
| Thiamine metabolism | Pyriethamine hydrobromide | Thiamine pyrophosphokinase | 34.48 | F |
| Arginine biosynthesis | H3B-120 | CPS1 | 15.4 | F |
| Glutathione metabolism | L-Buthionine-sulfoximine (BSO) | γ-glutamylcysteine synthetase | 221.8 | C |
|  | N-acetyl-L-methionine | Metabolite that decreases the hepatic glutathione level | 3138 | F |
| One carbon pool by folate | not tested | NA |  |  |
| Nicotinate and nicotinamide metabolism | not tested | NA |  |  |
| Vitamin B6 metabolism | Pyridoxine HCl | Competitive inhibitor for vitamin B6 | 71.22 | P |
| Biotin metabolism | not tested | NA |  |  |
| Pantothenate and CoA biosynthesis | not tested | NA |  |  |
| Folate biosynthesis | not tested | NA |  |  |

Of 20 tested drugs, 13 fully inhibited bacterial growth at non-cytotoxic drug concentrations (Figure 4a; Supplemental Figure 3), 6 fully inhibited bacterial growth but with a concomitant increase in host cytotoxicity at the minimum inhibitory concentration, 2 showed partial inhibition, and only 1 had no effect on bacterial growth (Figure 4a; Supplemental Figure 3). The standard pathogen-targeting antibiotic tetracycline (IC_50_ = 0.03 μM) served as a positive control (Figure 4a). Among the inhibitory drugs with no cytotoxic effect, 5 exhibited an IC_50_ < 5 μM (Figure 4a), constituting 23% of all inhibitors tested. These five were mycophenolate mofetil (IC_50_ = 1.13 μM) and pitavastatin (IC_50_ = 4.44 μM), which are FDA-approved, brequinar (IC_50_ = 0.86 μM) which has gone through phase II clinical trials, and 6-aminonicotinamide (IC_50_ = 1.16 μM) and roseoflavin (IC_50_ = 0.60 μM) which have not yet been tested for safety or efficacy in humans. Drug indication and pharmacokinetic information are provided in Supplemental Table 2. Importantly, both mycophenolate mofetil and brequinar have serum concentrations that exceed the IC_50_ concentrations derived here, making it likely that these drugs will be effective at the dosages that have been deemed safe for humans.

**Figure 4.**
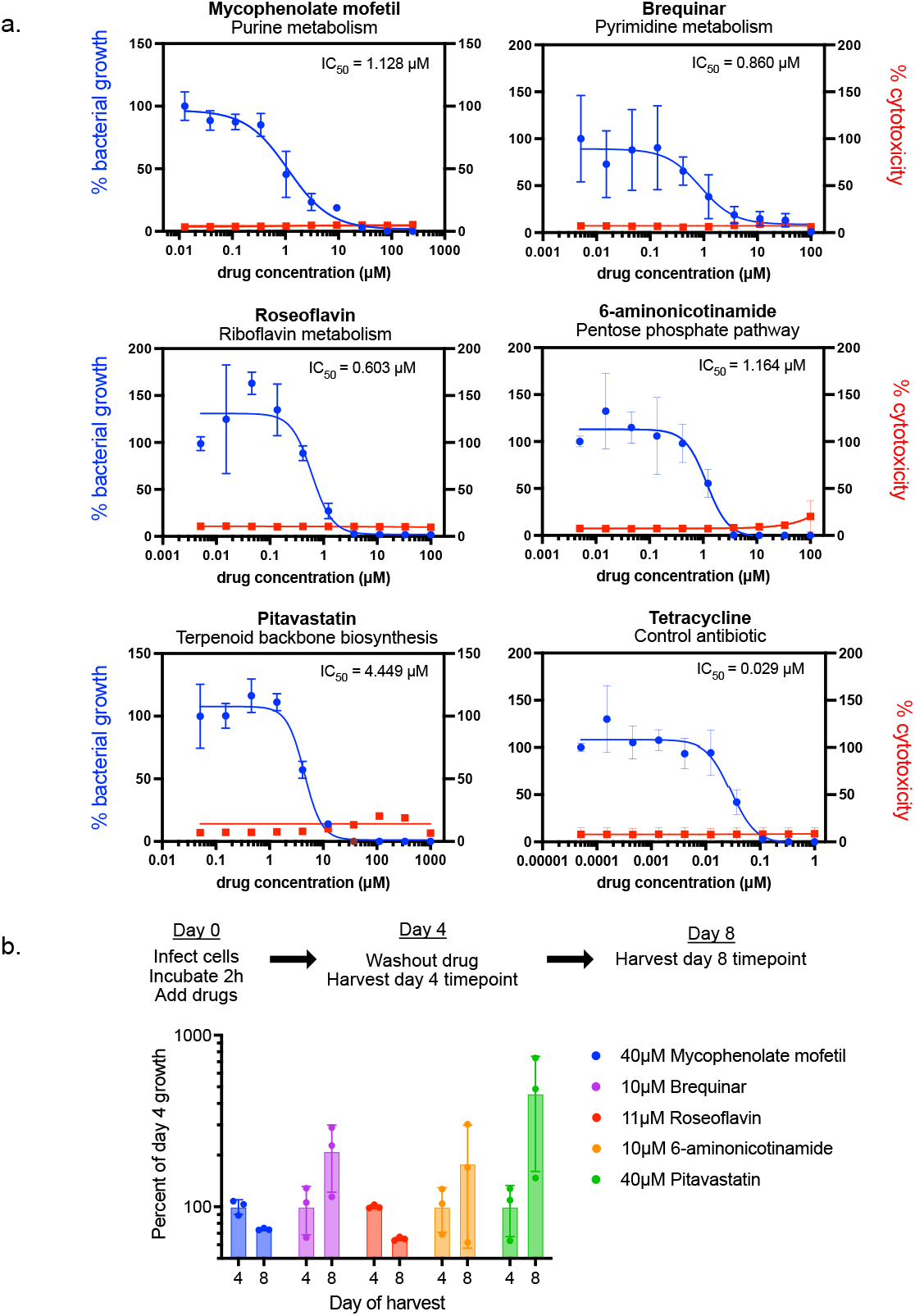
*In vitro* validation of host directed therapeutics against *Rickettsia parkeri* infections. a) Dose response curves for 5 host-directed therapeutics with IC_50_ < 5μM and minimal host cytotoxicity, with tetracycline as a control. b) Washout experiment to test the mode of killing of the inhibitors shown in panel A. Concentrations are listed in the figure legend and exceed the estimated minimum inhibitory concentration (MIC).

### Assessing mode-of-killing by lead compounds

To determine whether the mode of drug inhibition is bacteriostatic or bactericidal (the desired mode), we assayed the restoration of bacterial growth after discontinuing drug treatment. We focused on drugs with no cytotoxic effects and an IC_50_ < 5μM, using tetracycline as a control. Bacteria-infected cells were incubated for 4 days with a drug concentration that exceeds the predicted minimum inhibitory concentration (MIC), the lowest concentration at which bacterial growth is completely inhibited based on their respective dose response curves (Figure 4b; Supplemental Table 2). The drug media was then washed out and replaced with fresh media without drugs, and the cells were incubated for 4 additional days to allow any surviving bacteria to replicate.

Mycophenolate mofetil and roseoflavin (median fold change in bacterial numbers from day 4 to day 8 = 0.74 and 0.65, respectively) displayed activity consistent with a bactericidal mode of growth inhibition, as bacterial load continued to decrease even in the absence of the drug. In contrast, pitavastatin, brequinar, and 6-aminonicotinamide displayed activity consistent with a bacteriostatic mode of growth inhibition, as bacterial numbers increased 4.6, 2.1, and 1.8-fold from day 4 to day 8.

### Metabolomics of infected cells

To further validate our genomics-guided predictions of host metabolic dependency, we pursued a complementary approach and employed unbiased metabolomics to evaluate the global metabolic changes due to *Rickettsia* infection of host cells. We hypothesized that bacterial consumption of host metabolites would affect overall metabolite levels within the pathways identified in this study. To test this hypothesis, we compared global metabolite and lipid levels from infected and mock-infected A549 cells that were harvested 4 days after infection in five parallel replicates. Differences in the intracellular metabolite (Figure 5a) and lipid (Supplemental Figure 4) peak intensities between the infected and mock-infected conditions were visualized on volcano plots. We observed few significant changes in the lipid fraction except for several ether-linked phosphatidylglycerols, which are abundant components of bacterial membranes (Sohlenkamp and Geiger, 2016), and as expected were detected in the infected samples but completely absent in the mock-infected cells (Supplemental Figure 3). In contrast, we observed significant changes in metabolite levels, with 68 metabolites showing a decrease (log_2_ fold change <-0.5, *P* < 0.1) and only 10 metabolites showing an increase (log_2_ fold change >0.5, *P* < 0.1) in infected cells, consistent with our initial hypothesis of bacterial depletion of host metabolites (Figure 5a). Pathway enrichment analysis using Metaboanalyst 5.0, which estimates the importance of a given pathway as determined by the total changes in measured metabolites, identified 14 significantly altered pathways (pathway impact score > 0.1, FDR *P* < 0.1; Supplemental Table 3; Chong et al., 2018).

**Figure 5:**
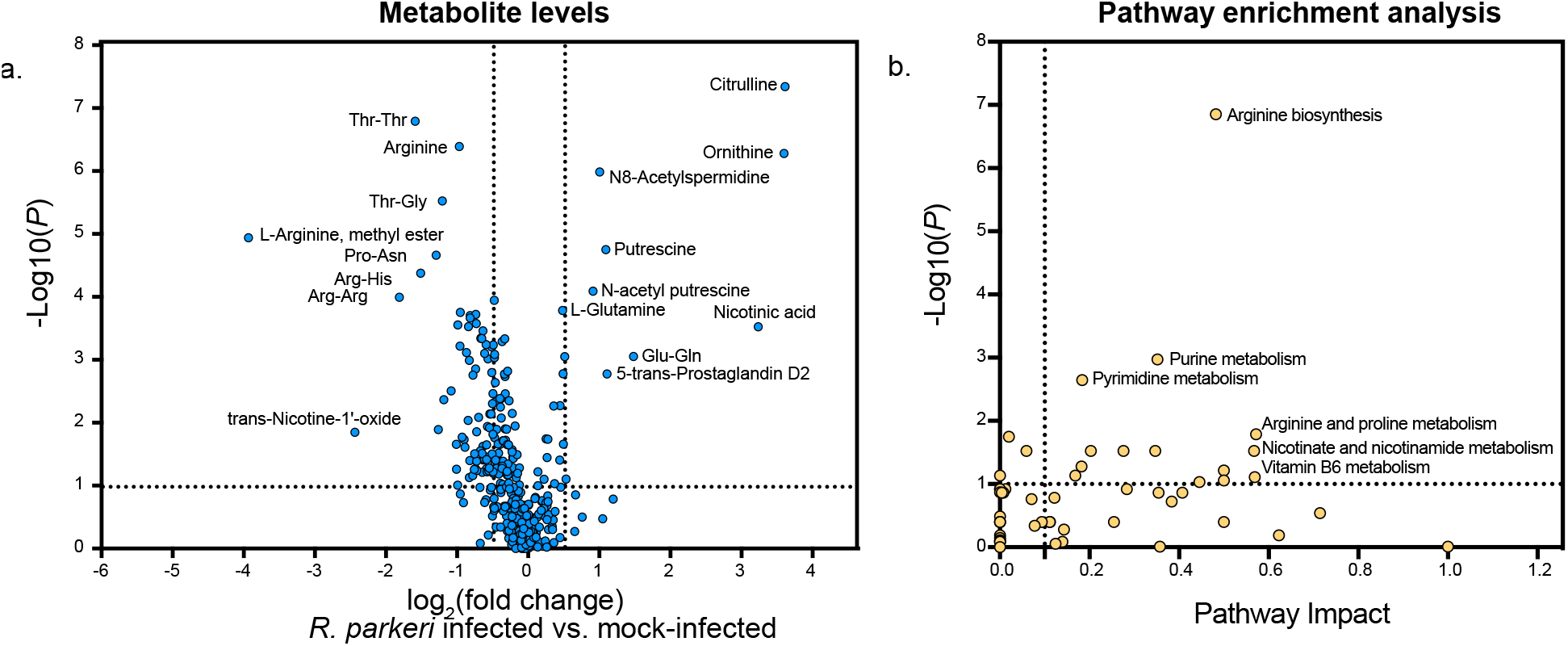
Metabolomics of *Rickettsia parkeri*-infected versus mock-infected A549 cells after 4 days of infection. a) Volcano plot of metabolite levels measured by mass spectrometry. b) Pathway enrichment analysis showing pathway impact scores versus false discovery rate *P* values.

To illustrate this pathogen-induced metabolic re-wiring of the host, we mapped three of the highest ranking pathways identified in the enrichment analysis, arginine biosynthesis (Pathway Enrichment rank = 1; PoMeLo rank = 14), purine metabolism (Pathway Enrichment rank = 2, PoMeLo rank = 1), and pyrimidine metabolism (Pathway Enrichment rank = 3, PoMeLo rank = 2), using the Kyoto Encyclopedia of Genes and Genomes (KEGG) pathway mapper tool (Supplemental Figures 5, 6, and 7, respectively; Kanehisa et al., 2022, 2016). Our PoMeLo pipeline would predict that the *Rickettsiaceae* acquires arginine, purines, and pyrimidines from the host.

The pathway showing the most change in the infected cells was arginine biosynthesis. The presence or absence of genome-encoded enzymes for a representative member of the *Rickettsiaceae (R. parkeri)* is diagrammed in Figure 6a, and a more detailed KEGG map is in Supplemental Figure 5. Our analysis found that arginine levels were decreased in infected versus mock-infected conditions (log_2_ fold change = -0.97, *P* = 4.08e-7). The large decrease in L-arginine methyl ester (log_2_ fold change = -3.93, *P* = 1.15e-5, Figure 5b), while not a direct intermediate, also indicated perturbation of the arginine metabolic pathway. Interestingly, we saw the largest metabolite increases in ornithine (log_2_ fold change = 3.60, *P* = 5.26e-7) and citrulline (log_2_ fold change = 3.61, *P =* 4.57e-8), suggesting either a block in the synthesis of downstream products such as arginine or a host feedback response to arginine depletion that led to a corresponding upregulation of the host *de novo* arginine biosynthesis pathway. In summary, the genomic data reveal that the key enzymes to convert intermediates in the arginine pathway to the final product, arginine, are absent in the *Rickettsiaceae*, and the metabolomics data support the idea that Rickettsia are siphoning arginine from the host.

**Figure 6:**
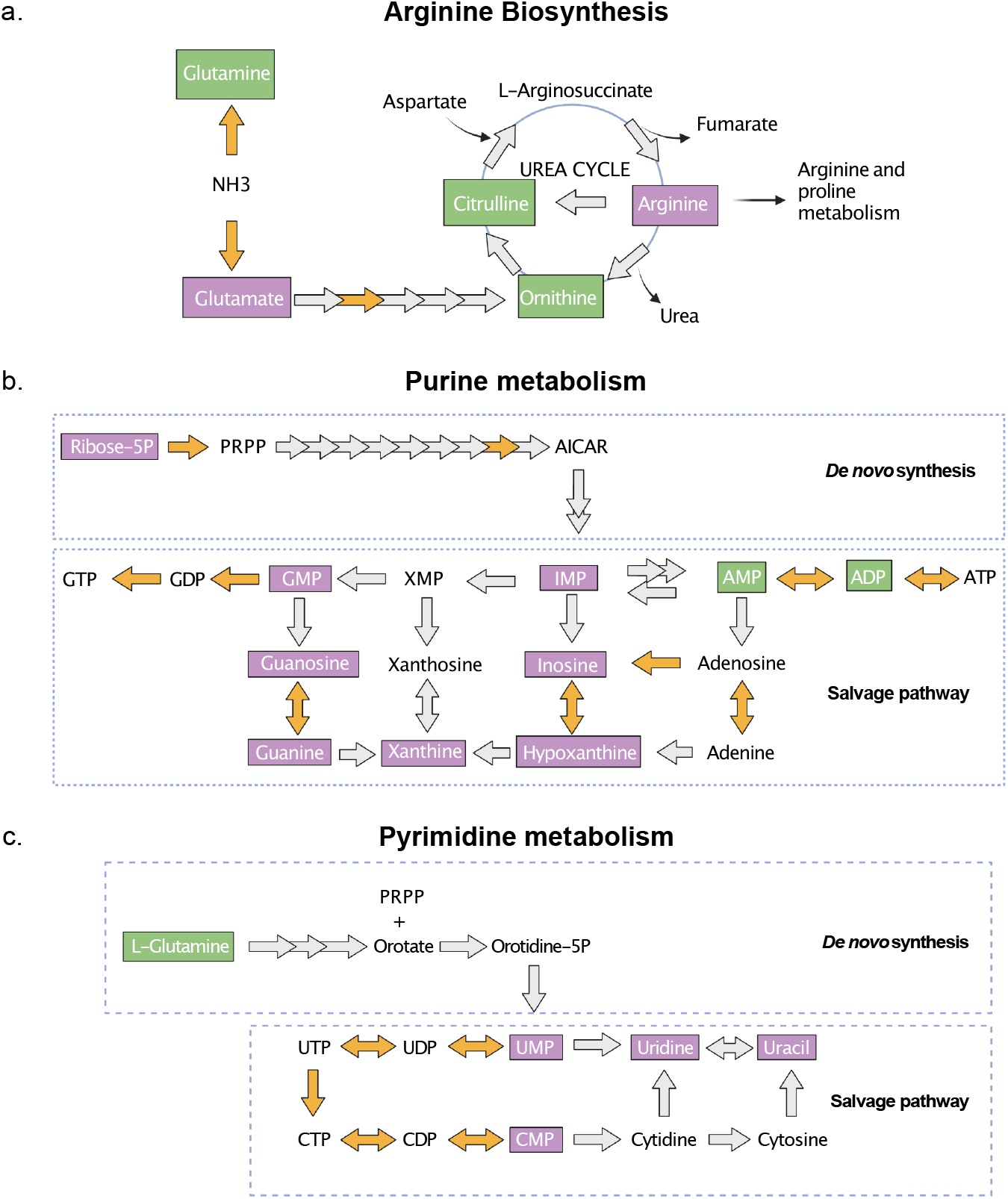
Metabolic pathway analysis of three top-scoring pathways from the metabolomics comparison of infected versus mock-infected samples. Schematics are shown for a) arginine biosynthesis, b) purine metabolism,, and c) pyrimidine metabolism (created using BioRender). Orange arrows signify enzyme presence and gray arrows signify enzyme absence in the *Rickettsia parkeri* genome, a proxy for the *Rickettsiaceae* target group. Colored boxes around individual metabolites signify either higher (purple) or lower (green) levels found in infectedversus mock-infected samples.

The purine metabolism pathway is split into a *de novo* biosynthesis pathway, and the salvage pathway (Figure 6b; Supplemental Figure 6). Mapping the genes encoded in the *Rickettsiaceae* revealed that the bacteria lack the ability to synthesize IMP from the *de novo* pathway (Figure 6b), unlike the related *Anaplasmataceae* (Supplemental Figure 6). Our metabolomic analysis revealed that most detected purine metabolites, including IMP, were low in infected cells (log_2_ fold change = -0.56, *P* = 0.61), consistent with their consumption by the bacteria. In contrast, elevated levels of AMP (log_2_ fold change = 0.27, *P* = 0.15) and ADP (log_2_ fold change = -0.02, *P =* 0.30) were observed, as expected given that *R. parkeri* expresses many enzymes that can convert imported ATP into ADP, AMP, and other intermediates in the salvage pathway (Atkinson and Winkler, 1985; Audia and Winkler, 2006).

The pyrimidine metabolism pathway is also split into a *de novo* pathway, where glutamine is converted into UMP, and the salvage pathway, where UMP and CMP can be converted into UTP and CTP. The *Rickettsiaceae* lack the ability to make UMP through the *de novo* pathway, unlike the *Anaplasmataceae* (Figure 6c; Supplemental Figure 7). However, in the salvage pathway, both the *Rickettsiaceae* and *Anaplasmataceae* encode enzymes that convert UMP and CMP into the final products, UTP and CTP. Consistent with this, we found that the pyrimidine intermediates UMP, CMP, uridine, and uracil were all depleted in infected cells. A previous biochemical study of *Rickettsia prowazekii* showed that the bacteria import host UMP as a precursor for the synthesis of UTP and CTP (Winkler et al., 1999). Altogether, the enzyme presence and metabolite levels indicate that the *Rickettsiaceae* cannot synthesize UMP and thus must consume host UMP for conversion of essential pyrimidines though the salvage pathway.

Intriguingly, of the 14 pathways identified as significantly altered (pathway impact > 0.1, FDR *P* < 0.1) in the metabolomics study, 11 (79%) were also identified in the top 20 pathways scored by PoMeLo (Figure 7). Two of the three remaining pathways that were not identified by the PoMeLo pipeline, the pentose and glucuronate interconversions and TCA cycle, had measurable PML scores (92 and 26, respectively) that did not reach the threshold of our top predicted pathways. Five of the nine pathways identified by PoMeLo but not by metabolomics (pentose phosphate pathway, cysteine and methionine metabolism, fructose biotin metabolism, and thiamine metabolism) were identified as altered by Metaboanalyst but did not pass the thresholds for significance. The extent of overlap between the two distinct but complementary approaches of metabolomics and genomic analyses further supports the idea that in the course of genome streamlining, these bacteria have jettisoned metabolic enzymes in favor of acquiring essential metabolites from the host. Therefore, limiting the host contribution to bacterial metabolism is a viable strategy to effectively disrupt or inhibit pathogen growth.

**Figure 7:**
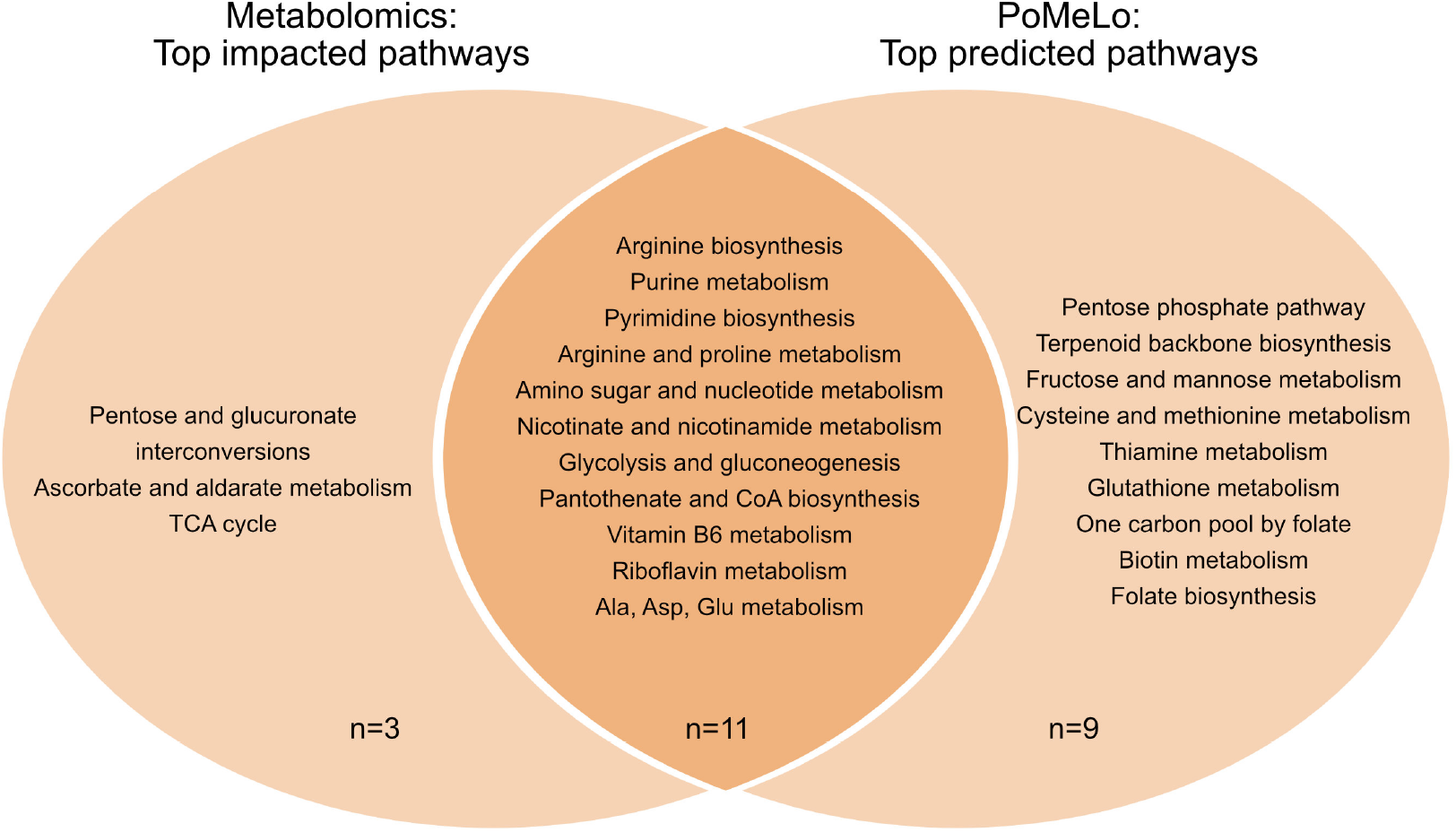
Venn diagram of significant metabolic pathways identified by unbiased metabolomics and the PoMeLo prediction pipeline, not including the 3 excluded pathways. Metabolomics pathways are filtered by impact factor > 0.1 and FDR P-value < 0.1.

## Discussion

We computationally identified 20 metabolic pathways that show significant gene loss in the *Rickettsiaceae* family but not in the closely related *Anaplasmataceae* family. To compensate for these incomplete pathogen-encoded pathways, obligate intracellular microbes must steal host metabolites to support their growth. Indeed, we showed that 59% of known inhibitors of host enzymes in these missing metabolic pathways strongly suppressed pathogen growth at non-cytotoxic concentrations. This high hit rate for inhibiting *in vitro* bacterial growth lends promise for further *in vivo* testing for host-directed therapeutics.

We also identified pathogen dependencies on host-generated metabolites by comparing overall metabolite levels in infected versus mock-infected samples. Strikingly, of 20 pathways identified by PoMeLo predictions and 14 pathways identified by unbiased metabolomics, 11 overlapped between the two distinct but complementary approaches. Thus, unbiased metabolomics is a promising approach to capture a large fraction of the metabolic pathways that pathogens co-opt from their host cells without the requirement of high-quality genome and protein annotations, and it would be especially useful for identifying therapeutics for emerging and novel pathogens.

Taking a host-directed therapeutic approach has considerable advantages over pathogen-targeting therapeutics. First, the wealth of pre-existing clinical data demonstrating the safety of existing drugs that target human metabolic processes can significantly expedite their FDA approval for a new indication (Ashburn and Thor, 2004). Second, the requirement for a drug to penetrate a pathogen cell is eliminated when its target occurs in the host cell. Third, while the rapid acquisition of drug resistance mutations is a major limitation for pathogen-targeting therapeutics (Hutchings et al., 2019), drug resistance is highly unlikely to occur with host-encoded targets. Finally, because our computational strategy relies on prior knowledge and predictions of the targets for host-directed therapeutics, it bypasses the costly and time-consuming target discovery and screening process.

Many of the drugs we identified in this study have gone through various stages of clinical development and therefore have available data from safety and efficacy studies. Existing pharmacokinetic profiles can help in understanding and choosing an appropriate dosage to be efficacious against pathogen growth while limiting host cytotoxicity. Our study identified five drugs, 23% of those tested, that inhibit bacterial growth with an IC_50_ < 5 μM. Of these, mycophenolate mofetil has the most promising properties: it is a potent (submicromolar IC_50_) FDA-approved compound with pharmacokinetic data that suggests its efficacy in humans, and our work shows that it not only inhibits growth but leads to bacterial cell death.

Although this study focuses on bacterial pathogens within the Alphaproteobacteria, genomic streamlining in pathogens is a well-documented phenomenon that occurs in diverse bacteria, eukaryotic parasites, and fungal pathogens (Moran, 2002; Poulin and Randhawa, 2015; Stajich, 2017). Our accompanying manuscript describing a systematic computational approach to compare bacterial genomes of target and non-target species will allow researchers to predict host dependencies from their pathogen of choice (Glascock, Waltari *et al* 2023), enabled by the abundance of well-annotated bacterial genomes available from data repositories such as NCBI and BV-BRC. In theory, this same type of analysis can be used for eukaryotic genomes, but the currently limited phylogenetic breadth of complete and well-annotated genomes for parasitic and fungal pathogens would likely result in incomplete predictions of host dependency. With the continual reduction of genome sequencing costs and faster tools for genome reconstruction and annotation, this barrier can be overcome as more genomes are added to central data repositories.

Furthermore, our complementary metabolomic approach was able to capture most of the same metabolic pathways as our PoMeLo predictions and could be employed for identifying metabolic dependency when insufficient genomic data is available.

Despite the promise of host-directed therapeutics for infectious diseases, some challenges remain to be considered. If pathogens hijack specific host metabolic pathways, will further inhibiting these host pathways during infection cause additional harm to the host? This scenario could be addressed by preclinical *in vivo* therapeutic studies in actively infected animals. Our work tests drug efficacy in a cell culture model, but the drug may target specific tissues that do not overlap with the pathogen’s tropism. In these cases, additional medicinal chemistry may be required to target the drug to the appropriate tissues.

We examined whether host-directed therapeutics show bactericidal or bacteriostatic effects on pathogenic bacteria and the differences found among the five best inhibitors can aid in selecting lead compounds. Understanding the mode of action of each drug is especially important when the target is the host cell and not the pathogen, as evidenced by the observed increase in bacterial load after removal of host-targeted bacteriostatic drugs. Although further safety and efficacy studies are required before host metabolic inhibition becomes a commonly utilized approach for host-directed therapeutics, the ability to make target predictions based solely on existing genomic information offers a tantalizing promise of transforming the field of novel antimicrobial drug discovery.

## Materials and Methods

### Reagents

FBPase inhibitor (18860), Lometrexol (18049), and Pyrithiamine hydrobromide (19526) were obtained from Cayman Chemical Company. N-acetyl-DL-methionine (A010025G) was obtained from Fisher Scientific. Sparfosic acid trisodium (HY-112732B) was obtained from MedChem Express. Lumiflavine (SC-224045), Roseoflavin (SC-208315C), and Tetrahydro-2-furoic acid (SC-253674) were obtained from Santa Cruz Biotechnology, Inc. AP-III-a4 hydrochloride (S7443-2mg), 3PO (S7639), Sodium dichloroacetate (S8615), and Telaglenastat (S7655) were obtained from Selleck Chemicals. Pitavastatin (SML2473-5MG), Pyridoxine HCl (P6280-25G), DL-Propargylglycine (P7888-1G), Tetracycline (T7660-5G), L-Buthionine-sulfoximine (B2515-500MG), H3B-120 (SML3007), 6-aminonicotinamide (A68203-1G), Mycophenolate mofetil (SML0284-10MG), Ribavirin (R9644-10MG), and Brequinar (5083210001) were obtained from Sigma Aldrich. CBR-470-1 (7018) was obtained from Tocris. Splash Lipidomix lipid internal standards were obtained from Avanti Polar Lipids (330707).

### PoMeLo computational pipeline

A more thorough methods section is a part of the accompanying manuscript by Glascock, Waltari *et al* 2023. Complete genomes of bacteria in the *Rickettsiaceae, Anaplasmataceae*, and representative Alphaproteobacteria marked as ‘good quality’ were downloaded from BV-BRC. A phylogenetic tree using BV-BRC’s Bacterial Genome Tree tool was generated with the following parameters: 100 genes, 10 allowed deletions, and no duplications.

Phylogenetically-weighted mean genome sizes were then computed across this tree. In this procedure, either actual genome sizes (if the branch is an external tip) or estimated mean genome sizes (if the branch is an internal node) of descendant branches of an internal node are summed and then divided by the number of descendant branches. This phylogenetically-weighted estimate is propagated from the tips to the root. Since members of the Rickettsiales have uniquely undergone genome reduction within Alphaproteobacteria, the root of the Alphaproteobacteria tree was assigned the highest estimated genome size of its children branches (*i*.*e*. the value computed for the common ancestor of the outgroup, “other Alphaproteobacteria”) rather than a phylogenetically-weighted mean.

After choosing the target and non-target groups for analysis, a list of the genome IDs for each group is loaded into the PoMeLo pipeline, which downloads the annotation files for each genome and loads all the information into a single large dataframe. Gene annotations for each metabolic pathway are then queried across all target and non-target genomes, and the total differences at the gene level for each pathway are calculated to produce a predicted metabolic loss (PML) score per pathway. All the pathways are then ranked in a final csv output file with the PML scores and accompanied by two types of heatmaps. One type shows the target vs. non-target group comparison of enzyme presence per pathway, and the other shows percent presence of each pathway in the target vs. non-target group. All output files from the *Rickettsiaceae* vs. *Anaplasmataceae* comparison can be found in the Github repository for PoMeLo: https://github.com/czbiohub-sf/pomelo.

### Cell culture and *R. parkeri* infections

Confluent low-passage human lung epithelial A549 cells were obtained from ATCC and grown at 37°C in 5% CO_2_ and in high-glucose Dulbecco’s modified Eagle’s medium (DMEM; Gibco/Life Technologies) supplemented with 10% fetal bovine serum (Omega Scientific). Aliquots of bacteria were frozen at -80°C, and each infection was initiated from a single thawed 5 μl aliquot of wild-type bacteria. *R. parkeri* was propagated by infecting cell monolayers of A549 cells with wild-type *R. parkeri* at a multiplicity of infection (MOI) of 0.05 and grown at 33°C in 5% CO_2_ in DMEM supplemented with 10% FBS for four days.

### Dose response curves

Each compound underwent a 10-point, three-fold serial dilution dose response assay performed in triplicate within the innermost wells of a 1 mL 96 deep well plate. For mock wells, the highest concentration of DMSO was added. The contents of the drug dilution plate were added to the confluent A549 cell monolayers of the 96-well tissue culture plates 2 h following initial *R. parkeri* infection. Plates were then incubated at 33°C in 5% CO_2_. At four days post-infection and drug treatment, genomic DNA was extracted (*Quick*-DNA 96 kit; Zymo Research) for qPCR. Mode-of-killing assays were performed in triplicate in the same fashion, except that half the drug-treated infected wells were harvested on day 4, while the other half was washed and resuspended in fresh DMEM and harvested on day 8.

### qPCR quantification of *R. parkeri* growth

For qPCR quantification, 5 μl of genomic DNA was used with Luna Universal qPCR Master Mix (NEB) and primers for the *R. parkeri* gene encoding the 17-kDa antigen. A 10-fold serial dilution of a plasmid containing a single copy of the *R. parkeri* 17-kDa gene was used to create a standard curve for calculating *Rickettsia* genome copy number.

### Lactate dehydrogenase (LDH) assays

For LDH assays, A549 human lung epithelial cells (CCL-185™; ATCC) were plated and grown to confluence in a 96-well plate in 100 μl of DMEM (Gibco) supplemented with 10% FBS (Omega Scientific). Plates were then treated with the appropriate concentration of drug for 2 h at 33°C in 5% CO_2_. Following a 4-day incubation, host cell cytotoxicity due to drug treatment was measured (Cytotox-One Homogeneous Membrane Integrity Assay #G7891; Promega). Plates were read at excitation of 560 nm and emission of 590 nm using a fluorescence microplate reader. The maximum LDH release was an average of four untreated wells (A1-A4). Each value was then divided by this average maximum LDH release and multiplied by 100 to obtain percent lysis. Each experiment was performed, and data were averaged between biological replicates.

### GraphPad Prism analysis

Ct values were imported into GraphPad Prism (Version 9.5.1 (528)) and formatted into an XY graph where technical replicates for each drug were entered along with desired concentrations. The analysis tab enabled transformation of x-axis concentrations to common logarithms, followed by a nonlinear regression curve fit and sigmoidal, 4PL model where X is log(concentration).

### Metabolomics

A549 cells were seeded into 10 × 6-well tissue culture treated plates the day before infection. On the day of infection, cells were either infected with an MOI of 0.05 in 1 mL of DMEM (5 replicates) or mock-infected with DMEM only (5 replicates). Plates were centrifuged at 300 × *g* for 5 minutes and placed in a 33°C in 5% CO_2_ for 4 days.

After 4 days, cells from each well of a 6-well plate were washed with 2 mL of 1x PBS, scraped, and collected into an 1.5 mL centrifuge tube. Samples were washed twice with 500 μl of 1x PBS before resuspending in 225 μl ice-cold methanol with 1.5% iSTD-SPLASH and freezing on dry ice. Samples were processed within 48 h. Sample preparation and analysis have been detailed previously (https://www.protocols.io/private/8962C900975C11ED8E800A58A9FEAC02) Analysis of metabolomics data was performed on MetaboAnalyst 5.0 (Chong et al., 2018) and GraphPad Prism (Version 9.5.1).

## Data availability

The PoMeLo pipeline is available for download at https://github.com/czbiohub-sf/pomelo and includes all the input and output files from this manuscript and we have used Zenodo to assign a DOI to the repository: 10.5281/zenodo.8172742. Raw metabolomics and lipidomics data have been deposited to metabolomicsworkbench.org study ID ST002747 http://dx.doi.org/10.21228/M8P43M.

## Acknowledgements

We are grateful to Sandra Schmid, Joseph DeRisi, Rebecca Lamason and Matthew Welch for critical reading and feedback for this manuscript. We would also like to thank Thomas Burke and members of CZ Biohub for thoughtful discussion. Wild-type *R. parkeri* strain was provided by Dr. Matthew Welch. This work was supported by the Chan Zuckerberg Biohub-San Francisco.

